# Defective influenza A virus RNA products mediate MAVS-dependent upregulation of human leukocyte antigen class I proteins

**DOI:** 10.1101/2020.01.30.928051

**Authors:** Mir Munir A. Rahim, Brendon D. Parsons, Emma L. Price, Patrick D. Slaine, Becca L. Chilvers, Gregory S. Seaton, Andrew Wight, Sayanti Dey, Shannen L. Grandy, Lauryn E. Anderson, Natalia Zamorano Cuervo, Nathalie Grandvaux, Marta M. Gaglia, Alyson A. Kelvin, Denys A. Khaperskyy, Craig McCormick, Andrew P. Makrigiannis

## Abstract

Influenza A virus (IAV) increases presentation of class I human leukocyte antigen (HLA) proteins that limit antiviral responses mediated by natural killer (NK) cells, but molecular mechanisms have not yet been fully elucidated. We observed that infection with A/Fort Monmouth/1/1947 (H1N1) IAV significantly increased presentation of HLA-B, -C and -E on lung epithelial cells. Virus entry was not sufficient to induce HLA upregulation, because UV-inactivated virus had no effect. We found that HLA upregulation was elicited by aberrant internally-deleted viral RNAs (vRNAs) known as mini viral RNAs (mvRNAs) and defective interfering RNAs (DI RNAs), which bind to retinoic acid-inducible gene-I (RIG-I) and initiate mitochondrial antiviral signaling (MAVS) protein-dependent antiviral interferon (IFN) responses. Indeed, MAVS was required for HLA upregulation in response to IAV infection or ectopic mvRNA/DI RNA expression. The effect was partially due to paracrine signalling, as we observed that IAV infection or mvRNA/DI RNA-expression stimulated production of IFN-β and IFN-λ1, and conditioned media from these cells elicited a modest increase in HLA surface levels in naïve epithelial cells. HLA upregulation in response to aberrant viral RNAs could be prevented by chemical blockade of IFN receptor signal transduction. While HLA upregulation would seem to be advantageous to the virus, it is kept in check by the viral non-structural 1 (NS1) protein; we determined that NS1 limits cell-intrinsic and paracrine mechanisms of HLA upregulation. Taken together, our findings indicate that aberrant IAV RNAs stimulate HLA presentation, which may aid viral evasion of innate immunity.

**IMPORTANCE:** Human leukocyte antigens (HLA) are cell surface proteins that regulate innate and adaptive immune responses to viral infection by engaging with receptors on immune cells. Many viruses have evolved ways to evade host immune responses by modulating HLA expression and/or processing. Here, we provide evidence that aberrant RNA products of influenza virus genome replication can trigger RIG-I/MAVS-dependent remodeling of the cell surface, increasing surface presentation of HLA proteins known to inhibit the activation of an immune cell known as a natural killer (NK) cell. While this HLA upregulation would seem to be advantageous to the virus, it is kept in check by the viral non-structural 1 (NS1) protein, which limits RIG-I activation and interferon production by the infected cell.

## INTRODUCTION

Influenza A viruses (IAV) infect human airway epithelial cells and trigger innate host defences that limit virus replication and spread (1). Human respiratory epithelial cells are equipped with pattern recognition receptors (PRRs) including toll-like receptors (TLRs) and retinoic acid inducible gene I (RIG-I)-like receptors (RLRs) that bind viral RNA and transduce signals to initiate production of interferons (IFNs) and pro-inflammatory cytokines. In endosomes, TLR3 binds double-stranded (ds) viral RNAs (vRNAs) and initiates a signalling cascade to activate the pro-inflammatory transcription factor NFκB (2). However, during IAV infection of epithelial cells the bulk of viral RNA ligands for PRRs are located in the nucleus and cytoplasm; here, RIG-I serves as the chief sensor for IAV RNA species that include panhandle structures generated by complementary base-pairing of 5’ and 3’ ends of viral RNAs (3–8). Following vRNA binding, RIG-I associates with the mitochondrial antiviral signalling (MAVS) adaptor protein on the surface of mitochondria (9) and peroxisomes (10, 11); subsequent MAVS oligomerization causes recruitment and activation of interferon regulator factor IRF3 and IRF7 and transcription of antiviral type I IFNs (IFN-α and IFN-β) and type III IFNs (IFN-λ1-3).

Both *in vitro* and *in vivo* studies have shown that during viral RNA transcription and replication, IAVs generate defective RNA products missing portions of the viral RNAs (12). These include defective interfering (DI) RNAs, which are ≥178 nt long subgenomic RNAs that can be incorporated into defective viral particles (13); mini-viral RNAs (mvRNA) that are similar in structure to DI RNAs, but are considerably shorter (~56-125 nt long) (14); and the 22-27 nt long small viral RNA (svRNA) corresponding to the 5′ end of vRNA (15). Both DI RNAs and mvRNAs retain panhandle structures with closely apposed 5’ and 3’-ends that are ligands for RIG-I, which initiates antiviral signal transduction. Defective viral RNAs are thought to limit productive viral replication and the pathogenic effects of infection in part by being triggers for innate immune responses. mvRNAs are potent inducers of type I IFN production, whereas svRNAs fail to trigger IFN responses (14). However, it is unknown precisely how these defective viral RNAs affect the recognition of IAV-infected cells by the immune system.

Among the immune effector cells recruited to the lungs within days after IAV infection are natural killer (NK) cells, which possess cytotoxic function against virus-infected cells (16, 17). NK cells, whose function is regulated by an array of activating and inhibitory receptors, have an important role in the control of IAV infection in mice (18, 19). The activating NKp44 and NKp46, as well as co-stimulatory 2B4 and NTB-4 receptors aid in recognition and killing of IAV-infected cells by binding hemagglutinin (HA) protein on their surface (20–22). In mice, NKp46-deficiency results in increased morbidity and mortality following IAV infection, demonstrating the importance of this NK cell receptor in the control of infection (23, 24). Because binding of NKp46 to viral HA protein is dependent on sialylation of the O-glycosylated residues of NKp46, IAV can counter this recognition by cleaving the receptor sialic acids using the viral neuraminidase (NA) (25, 26). IAV can also circumvent NK cell-mediated antiviral responses by increasing the expression of inhibitory ligands, such as the class I human leukocyte antigen (HLA), also known as the human class I major histocompatibility complex (class I MHC), on the surface of infected cells. Class I HLA molecules are recognized by the human killer-cell immunoglobulin-like receptors (KIRs) on NK cells (27). Increased binding of inhibitory KIRs to class I HLA proteins on IAV-infected cells have been shown to inhibit NK cell function (28). Previously, we demonstrated that IAV infection in mice is associated with increased expression of mouse class I MHC on lung epithelial cells (29). On mouse NK cells the functional analogues of KIRs are inhibitory Ly49 receptors; we observed that disruption of inhibitory Ly49:class I MHC interactions improved survival of IAV-infected mice. Our study demonstrated that upregulation of class I MHC helps IAV evade NK cell-mediated immune responses, but the mechanism by which class I MHC is upregulated during IAV infection is not fully understood.

NK cell receptors bind to cognate ligands on the surface of infected cells and integrate activating and inhibitory signals that dictate the extent of NK cell activation (30). Knowing this, we initiated the current study to better understand how IAV infection affects the expression of ligands for NK cell receptors on the surface of infected epithelial cells. In-depth bioinformatic analysis of publicly available gene expression datasets revealed that IAV infection modulates the expression of a wide array of NK cell ligands, most notably, class I HLA genes that were consistently upregulated across many *in vitro* infection studies that employed different IAV strains and epithelial cell models. We complemented these findings using an A549 lung epithelial cell infection model. We observed significantly increased presentation of class I HLA and non-classical HLA-E on A/Fort Monmouth/1/1947 (H1N1) IAV-infected A549 cells. HLA upregulation was dependent on post-entry steps in replication because UV-inactivated virus had no effect. Specifically, we showed that IAV mvRNAs and DI RNAs are sufficient to increase HLA expression in the absence of infection. MAVS was required for HLA upregulation in response to IAV infection or ectopic mvRNA/DI RNA expression. IAV infection or ectopic mvRNA/DI RNA-expression stimulated production of IFN-β and IFN-λ, and conditioned media from these cells elicited modest increases in HLA presentation from naïve epithelial cells. Using the Janus kinase inhibitor Ruxolitinib (Rux) we demonstrated that signaling downstream of IFN receptors through Jak1 plays a major role in HLA upregulation triggered by IAV replication intermediates. Finally, we determined that IAV NS1 limits cell-intrinsic and paracrine mechanisms of HLA upregulation. Our data indicates that aberrant IAV mvRNAs and DI RNAs stimulate HLA presentation, which may aid viral evasion of immune surveillance.

## RESULTS

### Influenza A virus infection alters cell surface expression of ligands for NK cell receptors

NK cells control immune responses to IAV infection *in vivo* (18). NK cell receptors bind to cognate ligands on the surface of infected cells and integrate activating and inhibitory signals that dictate the extent of NK cell engagement (30). We performed an in-depth bioinformatic analysis of publicly available gene expression datasets (Table S1) to better understand how expression of known NK cell ligands is modulated by IAV infection *in vitro*. By focusing on datasets from *in vitro* IAV infections of standard epithelial cell models including primary human lung epithelial cells and alveolar adenocarcinoma A549 cells, we learned that expression of most known ligands for NK cell receptors is altered during IAV infection (Fig. 1A). In particular, there was a consistent trend of upregulation of HLA transcripts in multiple epithelial cell lines in response to infection by diverse IAV strains. These included the HLA-A, -B, -C, and -E proteins that present peptides to immune cells and bind inhibitory receptors on NK cells, as well as HLA-F, which binds to KIR receptors with context-dependent activating and inhibitory properties.

**Figure 1.**
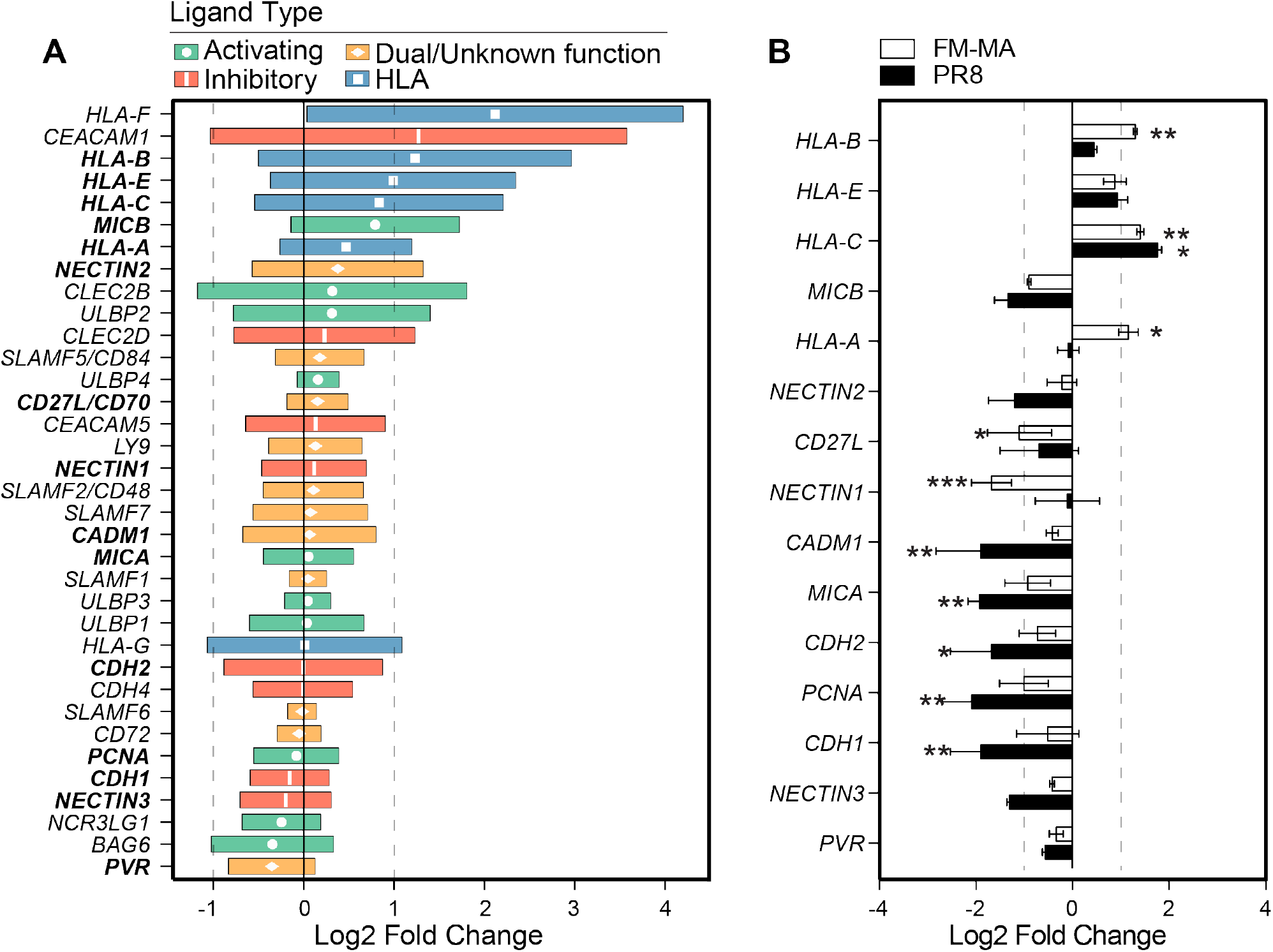
IAV infection of epithelial cells increases class I HLA gene expression. (A) Expression of NK cell ligands from 18 publicly available gene expression datasets from *in vitro* IAV infection of A549 cells and primary human lung cells. NK ligands are classified as activating (green), ambiguous function (orange) and nhibitory (red). Class I HLA proteins are indicated in blue. Data is presented as the log_2_ fold change relative o uninfected controls for each dataset; median values with interquartile range (IQR) are shown. Vertical dashed lines indicate 2-fold change thresholds. B) A549 cells were infected with PR8, FM-MA or mock-nfected for 17 h and RNA was harvested for RT-qPCR. Relative expression of NK cell ligands was expressed as log_2_ fold change relative to mock-infected controls. Vertical dashed lines indicate 2-fold change hresholds. N=3; * p<0.05, ** p<0.01, *** p<0.001.

To confirm reports of NK cell ligand modulation, we infected A549 cells with the A/Puerto Rico/8/1934 strain (PR8) at a MOI=1 for 16 h, at which point RNA was harvested and analyzed by RT-qPCR, which revealed statistically significant increases in HLA-C and significant decreases in MICA, MICB, NECTIN3, CADM1, CDH1, CDH2 and PCNA in PR8-infected A549 cells (Fig. 1B). By contrast, infection of A549 cells with the mouse-adapted A/Fort Monmouth/1/1947 (FM-MA) strain that we previously utilized to study NK cell responses to IAV infection in mice (29), caused significant increases in steady-state mRNA levels of HLA-A, -B, and -C, without causing statistically significant decreases in other NK cell ligands.

To determine whether changes in NK cell ligand mRNA levels led to corresponding changes in surface presentation of proteins, we infected A549 cells with PR8 or FM-MA and analyzed cell surface expression of NK cell ligands by flow cytometry (Fig. 2). We observed significant upregulation of HLA-A/B/C on the surface of PR8 and FM-MA infected cells (Fig. 2A) which correlated with our RT-qPCR data (Fig. 1B). When measured individually HLA-B, -C and -E were significantly upregulated by FM-MA infection, whereas PR8 infection elicited modest increases in HLA-B only, which did not achieve statistical significance (Fig. 2A). MICA/B ligands for the activating NKG2D receptor were differentially regulated by infection; PR8 infection had no effect on cell surface levels of MICA/B, whereas FM-MA infection caused a modest downregulation that agreed with our RT-qPCR data (Fig. 2A and Fig. 1B). There was a modest but statistically significant upregulation of CD155/PVR and downregulation of CD113/NECTIN3 in FM-MA infected cells (Fig. 2B). Downregulation of CD113/NECTIN3 was consistent with our RT-qPCR data (Fig. 1B) and bioinformatics analysis (Fig. 1A). Taken together, our bioinformatic analysis of published gene expression datasets, combined with our own RT-qPCR and surface staining experiments, clearly demonstrate that IAV infection alters the expression of NK cell ligands, and that the most striking and consistent finding is increased surface presentation of class I HLA proteins, in agreement with previous studies (28, 29).

**Figure 2.**
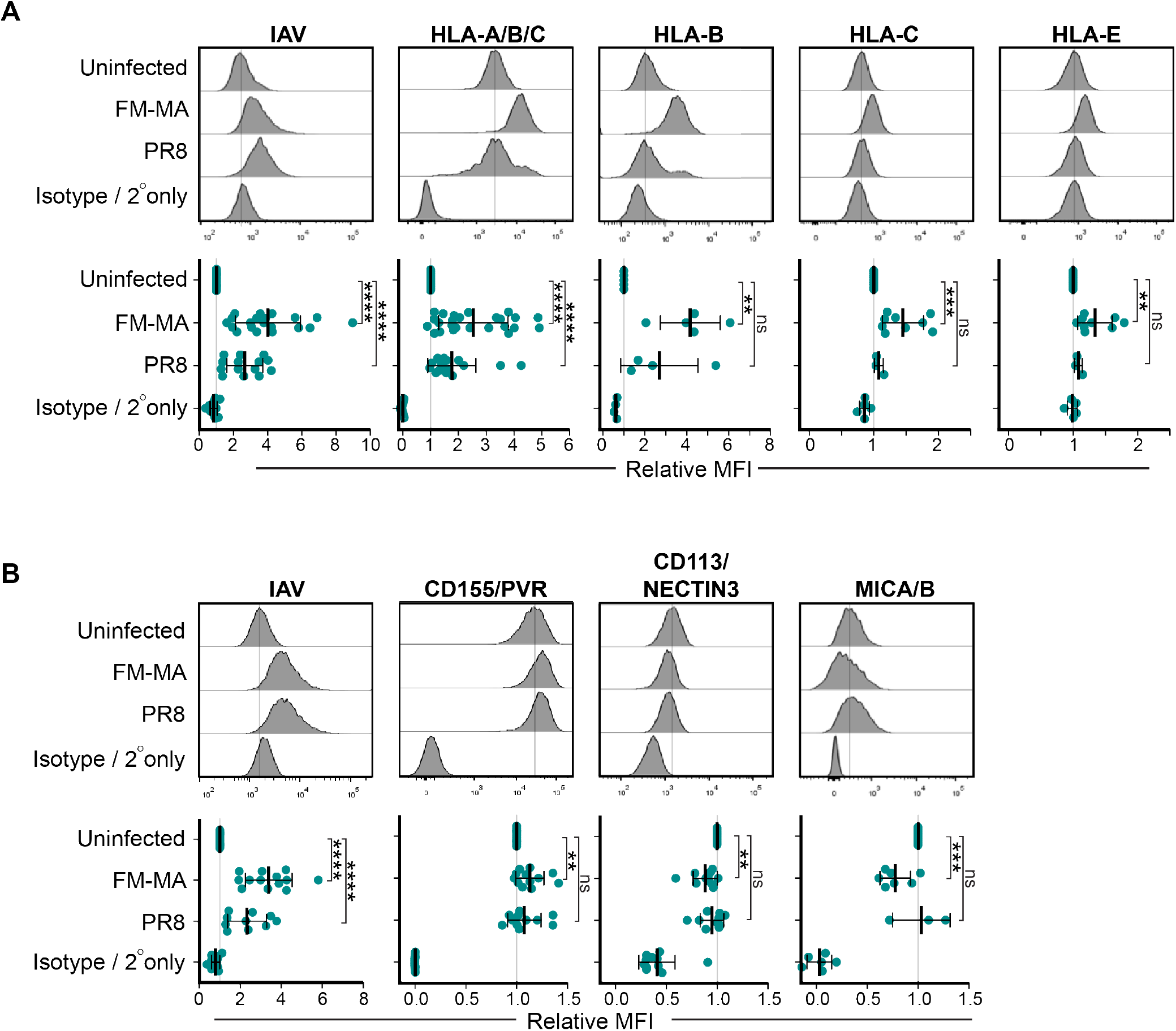
IAV infection alters cell surface expression of ligands for NK cell receptors. A549 cells were infected with FM-MA or PR8 at MOI=1. At 17 h, cells were fixed and immunostained to determine cell surface levels of NK cell ligands; cells were subsequently permeabilized for immunostaining of intracellular IAV proteins. (A) Flow cytometry analysis of cells immunostained with a pan-HLA-A/B/C antibody, or antibodies specific for class I HLA proteins HLA-B, HLA-C or HLA-E or isotype antibody controls. (B) Flow cytometry analysis for cells immunostained with antibodies to detect ligands for activating NK cell receptors; CD155/PVR, CD113/NECTIN3 and MICA/B or isotype antibody controls. Representative histograms (top panels) show results of a single experiment; vertical line indicates expression level of target in uninfected cells. Bottom panels show Mean Fluorescence Intensity (MFI) relative to uninfected cells. Each data point represents an independent experiment. Means and ±SD are shown. * p<0.05, ** p<0.01, *** p<0.001, **** p<0.0001.

### Defective vRNAs increase surface HLA presentation in a MAVS-dependent manner

To determine whether HLA upregulation was a consequence of IAV entry or later steps in viral replication we infected A549 cells with UV-inactivated or control FM-MA virus and measured cell surface HLA levels by flow cytometry using a pan-HLA-A/B/C antibody or an HLA-B-specific antibody. UV treatment damages viral RNA and prevents transcription and replication of the viral genome (31). We observed that, unlike infectious virus that increased cell surface HLA as expected, UV-inactivated inoculum had no effect (Fig. 3A).

**Figure 3.**
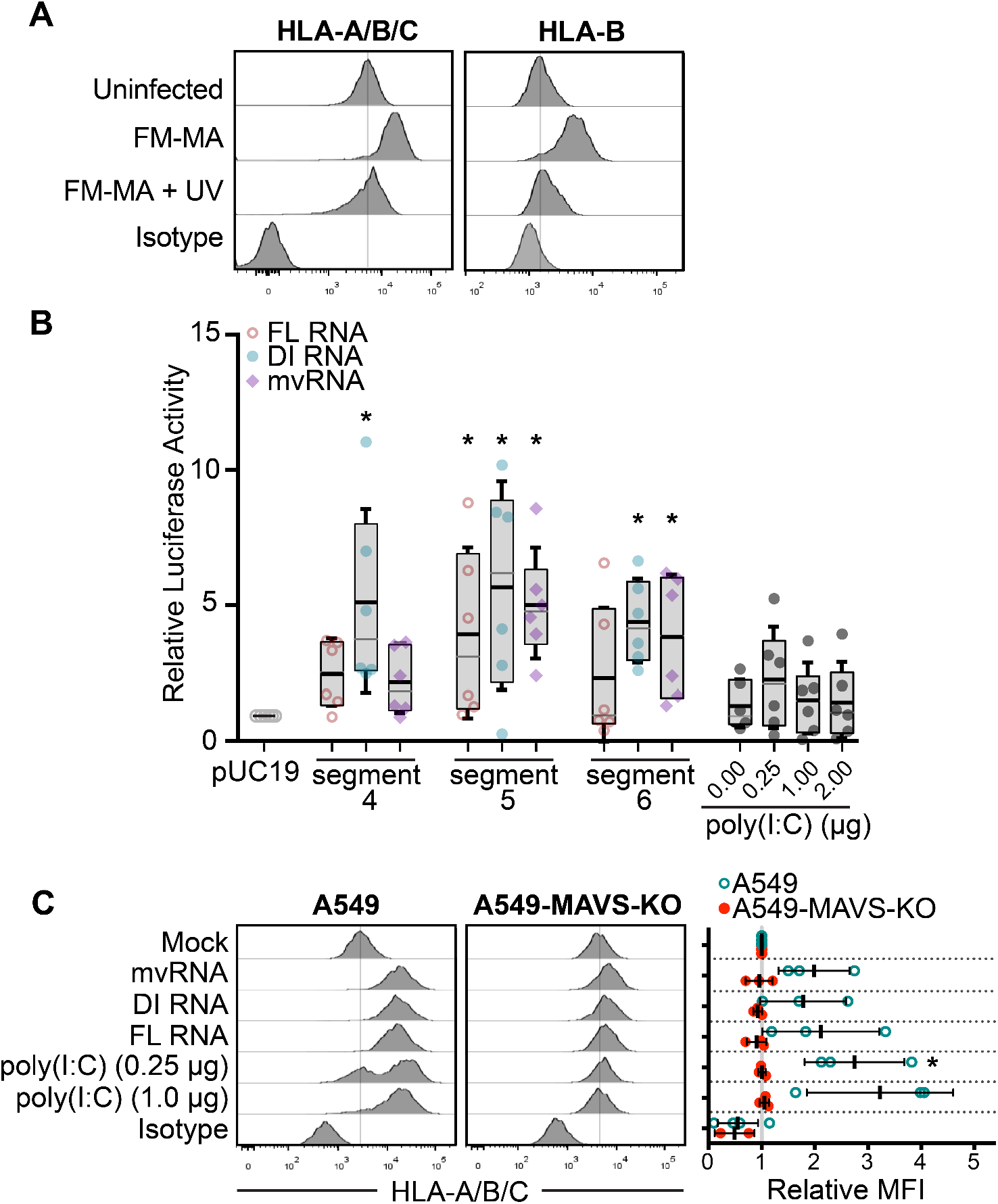
Defective viral RNAs increase surface HLA presentation in a MAVS-dependent manner. (A) FM-MA inoculum was exposed to UV light prior to infection of A549 cells at MOI=1. At 17 hpi, cells were fixed and immunostained with a pan-anti-HLA-A/B/C antibody or an anti-HLA-B antibody and processed for flow cytometry. Vertical line indicates HLA expression level in uninfected cells. Representative data from one out of two independent experiments is shown. (B) A549 cells were transfected with full length (FL) vRNA, defective interfering (DI) vRNA or mini-viral RNA (mvRNA) minireplicons derived from the ndicated genome segments. An ISRE-driven firefly luciferase reporter plasmid was co-transfected to measure IFN signaling, along with a Renilla luciferase plasmid that served as normalization control. Poly(I:C) and empty pUC19 plasmid served as positive and negative controls, respectively. Firefly luciferase activity was normalized to Renilla luciferase control for each sample, and data was expressed as fold change compared to pUC19 plasmid transfection (n=6, *p<0.05; IQR Boxes and SD whiskers are shown). (C) A549 cells or A549 MAVS-KO cells were transfected with IAV minireplicon expressing defective vRNAs from genome segment 5, as in (B), and analyzed by flow cytometry at 48 h post-transfection via surface mmunostaining with a pan-anti-HLA-A/B/C antibody (n=3). Histograms from a representative experiment are shown on the left; vertical line indicates expression level of target in uninfected cells. On the right, relative MFI values from at least 3 independent experiments are shown.

During replication the IAV RdRp frequently generates defective RNA products including DI RNAs (12) and smaller mvRNAs (14). Like intact full-length vRNAs, DI RNAs bind to the viral nucleoprotein (NP) and assemble into viral ribonucleoprotein (vRNP) structures that limit RIG-I binding (13). By contrast, mvRNAs do not bind to NP and are thought to be primary RIG-I agonists (14). One consequence of IFN signal transduction is increased cell surface HLA presentation (32). Because UV-inactivated IAV was unable to increase HLA surface presentation, we reasoned that increased HLA gene expression could be triggered by innate immune responses activated by defective RNAs produced during viral replication. To test the ability of defective RNAs to induce IFN signaling in our system, we used an IFN-β-responsive luciferase reporter driven by an interferon-stimulated response element (ISRE) promoter element. We observed that transfection of A549 cells with constructs bearing mvRNAs, DI RNAs or full-length vRNAs substantially induced ISRE-luciferase reporter activity (Fig. 3B). Interestingly, DI RNAs from genome segment 4 strongly activated ISRE-luciferase activity, whereas full-length vRNA or mvRNAs from the same segment had a moderate effect. By contrast, all three RNA species derived from genome segment 5 activated the ISRE-luciferase reporter to a similar extent. Notably, the virus-derived RNA species all potentiated stronger ISRE-luciferase reporter activity compared to poly(I:C), a known inducer of type I IFN in transfected A549 cells. Thus, in the A549 cell line used extensively in this study, diverse viral RNA species can elicit IFN signaling.

Because many viruses selectively modulate HLA presentation to disrupt antiviral immune responses (33, 34) we wondered whether defective IAV vRNAs might affect HLA presentation. To test this directly, we transfected A549 cells with constructs encoding the IAV mini-replicon system bearing mvRNA, DI RNA or full-length vRNA species, and measured surface levels of HLA. Control cells transfected with poly(I:C) showed strong dose-dependent upregulation of surface HLA-A/B/C over a 48 h period (Fig. 3C). Cells expressing viral RNAs from segment 5 likewise displayed strong surface HLA staining, indicating that they are sufficient to increase cell surface HLA in the absence of infection, in agreement with their ability to stimulate the ISRE-luciferase reporter (Fig. 3B).

RIG-I binds to IAV RNA panhandle structures and assembles with the MAVS adaptor on the surface of mitochondria and peroxisomes to drive antiviral signal transduction and IFN production (9, 10). To test whether the RIG-I/MAVS axis was involved in HLA upregulation in our system, we measured surface HLA-A/B/C expression in parallel in transfected MAVS-deficient A549 cells (A549-MAVS-KO) (details of construction and validation of A549-MAVS-KO cells in Fig. S1). Class I HLA levels on the MAVS-KO cells were unaffected by transfection with mvRNA, DI RNA or full-length vRNA constructs (Fig. 3C). This indicates that HLA upregulation in IAV-infected cells results from activation of the RIG-I/MAVS pathway that recognizes viral replication intermediates.

### Class I HLA upregulation in IAV-infected cells is MAVS-dependent

Having established that the RIG/MAVS axis is involved in HLA upregulation in response to ectopic expression of viral replication intermediates, we next addressed the role of RIG-I/MAVS in authentic IAV infection. A549 cells or A549-MAVS-KO cells were infected with FM-MA virus and class I HLA expression was measured by flow cytometry. At 17 hpi, class I HLA-A, -B and -C mRNA levels were significantly increased in infected WT A549 cells compared to mock-infected control cells but did not increase in MAVS-KO cells (Fig. 4A). Expression of other components of the antigen processing and presentation machinery including β2 microglobulin (B2M), transporter associated with antigen processing (TAP1) and proteasome subunit beta 8 (PSMB8) was significantly increased in A549 cells by 17 hpi, but not in MAVS-deficient cells (Fig. 4B). Surface class I HLA levels were largely unchanged in the early stages of infection, with moderate increases first measured at 12 hpi and increased further by 17 hpi (Figs. 4C, 4D). Cell surface levels of the non-classical HLA-E also increased on A549 cells over the infection time-course. By contrast, cell surface levels of these classical and non-classical HLA proteins remained unchanged in the A549-MAVS-KO cells throughout the time-course, despite robust accumulation of viral proteins indicative of progression of the infectious cycle (Fig. 4C, 4D). Together, these findings indicate that MAVS is required for IAV-induced HLA upregulation on the surface of infected A549 cells.

**Figure 4.**
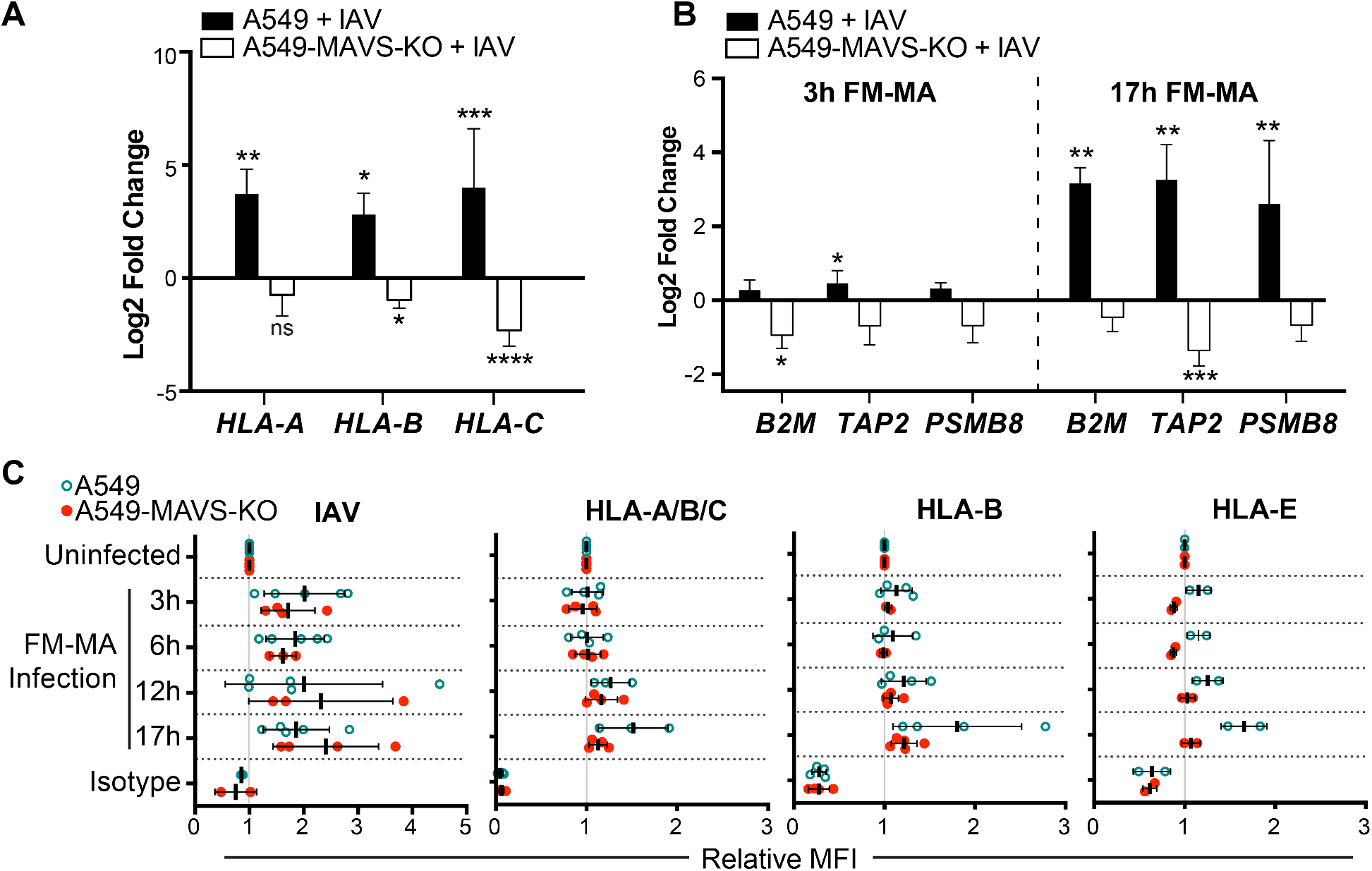
Class I HLA upregulation in IAV-infected cells is MAVS-dependent. A549 cells or A549-MAVS-KO cells were infected with FM-MA at an MOI=1. RNA was harvested for RT-qPCR at 3 hpi or 17 hpi. (A) Relative fold change in HLA-A, -B and -C transcript levels in A549 cells or A549-MAVS-KO cells at 17 hpi (n=3). (B) Relative fold change in B2M, TAP and PSMB8 transcript levels in A549 and A549 MAVS-KO cells at 3 hpi or 17 hpi (n=3). (C) Relative MFI of cell surface HLA proteins in FM-MA infected A549 cells and A549-MAVS-KO cells at indicated times, relative to uninfected controls.

### Defective IAV RNAs elicit cell-intrinsic and paracrine upregulation of class I HLA proteins

Because signaling downstream of type I IFN receptors increases class I HLA expression (32), we investigated the contribution of soluble factors to HLA expression in IAV infected cells. We infected A549 cells with FM-MA for 17 h and collected cell supernatants, which were UV-treated to inactivate virions prior to incubation with naïve A549 cells for an additional 17 h. Donor and recipient A549 cells were stained with anti-HLA-A/B/C or anti-HLA-B antiserum and analyzed by flow cytometry. We observed marked increases in surface class I HLA proteins on IAV-infected A549 cells as before, compared to moderate increases on cells incubated with UV-treated conditioned medium (Figs. 5A and B). Staining cells with anti-IAV antiserum confirmed that the UV-treatment of culture supernatants inactivated virions and prevented subsequent infection of naïve A549 cells, mitigating concerns of residual infection in these experiments (Fig. 5B). Incubating naïve A549 cells with culture supernatants collected from cells expressing IAV mvRNAs yielded a similar result, with strong significant increases in class I HLA protein levels on the donor cells compared to relatively modest increases on the cells that received the conditioned medium (Fig. 5C). Together, these findings indicate that class I HLA can indeed be upregulated on epithelial cells in a paracrine manner in response to infection, but this effect is weaker than the cell-intrinsic class I HLA upregulation on the infected cell.

**Figure 5.**
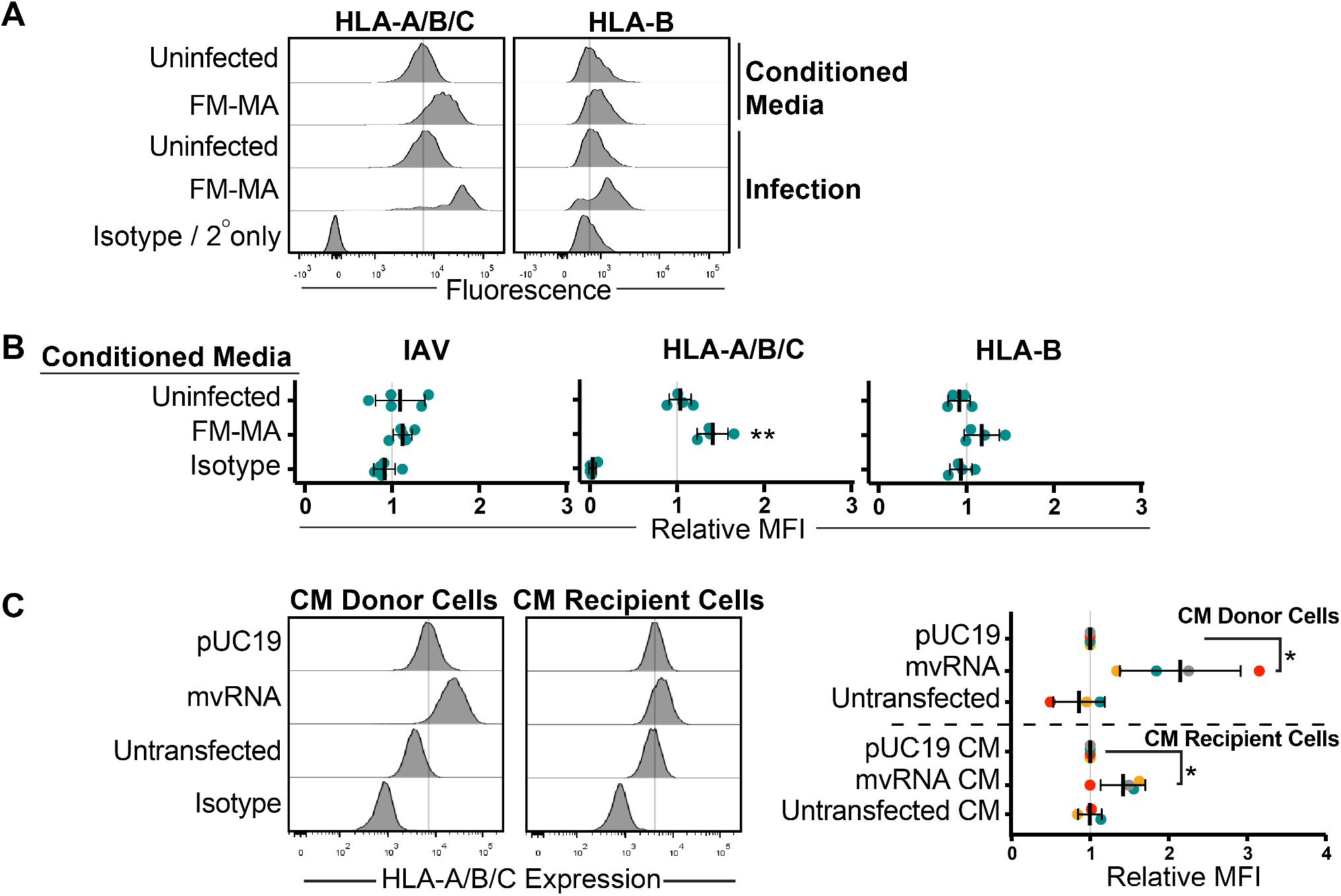
Defective IAV RNAs elicit cell-intrinsic and paracrine upregulation of class I HLA proteins. A549 cells were treated with conditioned medium containing UV-inactivated culture supernatant from FM-MA-infected cells. Surface HLA levels on recipient cells (17 h post-treatment) and infected donor cells (17 hpi) were determined by flow cytometry. Histograms from a representative experiment are shown. Vertical dashed-line indicates expression level in uninfected cells. (B) MFI of cell surface HLA proteins on recipient cells from (A) relative to cells treated with conditioned media from mock-infected cells. Each data point represents an independent experiment. (C) A549 cells were treated with conditioned medium from cells ransfected with IAV minireplicon expressing defective vRNAs from genome segment 5 or from control untransfected cells or pUC19 vector-transfected cells. After 24 h, cells were fixed and immunostained with a pan-anti-HLA-A/B/C antibody (n=3). Histograms from a representative experiment are shown on the left; vertical line indicates expression level of target in uninfected cells. On the right, relative MFI values from at east 3 independent experiments are shown (*p<0.05).

### HLA upregulation in response to defective IAV RNAs is dependent on IFN signaling

IAV infection induces production of type I and type III IFN proteins by the infected cell that orchestrate autocrine and paracrine anti-viral responses (1, 5). Compared to uninfected A549 cells, infection with FM-MA induced MAVS-dependent expression of *IFN-β* and *IFN-λ1* genes as early as 3 h post-infection, which increased to 50-fold and 150-fold, respectively, by 17 h post-infection (Fig. 6A). To confirm that type I and type III IFNs can induce HLA upregulation in our system, we treated A549 cells with IFN-β, IFN-λ1 or IFN-λ2, and compared HLA expression in IFN-treated and untreated cells. Both RT-qPCR and flow cytometry analysis showed that IFN-β was the most potent inducer of class I HLA mRNA and protein expression in A549 cells; HLA-A, HLA-B and HLA-C mRNAs accumulated in IFN-β-treated cells by 12 h post-treatment (Fig. 6B), which was reflected in increased HLA-A/B/C cell surface staining (Fig. 6C). By contrast, 12 h treatment with IFN-λ1 potently increased HLA-A mRNA levels, but not HLA-B and -C mRNA levels (Fig. 6B). Overall, IFN-β was a much more potent inducer of HLA in our system compared to IFN-λ1 and IFN-λ2.

**Figure 6.**
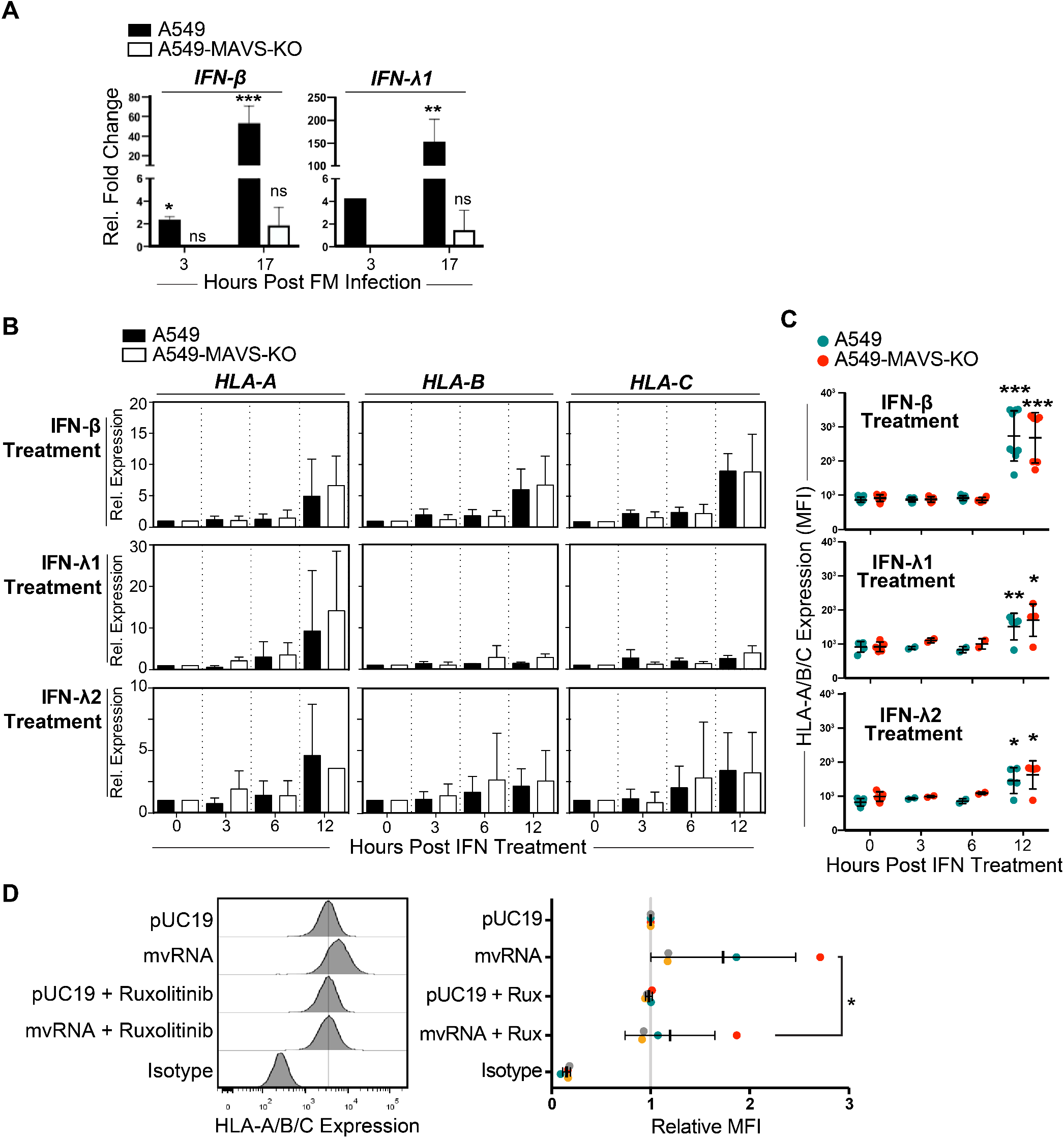
HLA upregulation in response to defective IAV RNAs is dependent on IFN signaling. (A) A549 cells or A549-MAVS-KO cells were infected with FM-MA for 17 h and relative levels of IFN-β and IFN-λ1 transcripts compared to uninfected controls were analyzed by RT-qPCR (n=3). (B) A549 cells or A549-MAVS-KO cells were treated with recombinant IFN-β, IFN-λ1 or IFN-λ2 and RNA was harvested over a 12 h time course. Relative expression of HLA-A, -B and –C transcripts was analyzed RT-qPCR. (C) Surface expression of HLA-A/B/C was determined by immunostaining and flow cytometry of cells harvested over the time course of IFN treatment described in (B) (n=3). (D) Analysis of HLA surface expression on A549 cells transfected with IAV minireplicon expressing defective vRNAs from genome segment 5 or from control pUC19 vector-transfected cells. Immediately after transfection, cells were treated with Ruxolitinib (Rux) or mock-treated. * p<0.05, ** p<0.01, *** p<0.001.

Autocrine and paracrine type I and III IFN signaling is mediated by IFN receptor signal transduction via downstream non-receptor tyrosine kinases, Jak1, Jak2 and Tyk2 (35–39). To directly test if IFN receptor signalling plays a role in HLA upregulation, we treated A549 cells with the Jak1 inhibitor Rux. In A549 cells transfected with mvRNA-expressing minireplicon, we observed that upregulation of HLA-A/B/C on the cell surface was inhibited by Rux treatment (Fig. 6D). In control pUC19-transfected cells, Rux had no effect on HLA levels. Together, these data clearly indicate that signaling downstream of IFN receptors through Jak1 plays a major role in HLA upregulation triggered by IAV replication intermediates.

### NS1 protein limits cell-intrinsic and paracrine upregulation of class I HLA proteins

In many experiments FM-MA infections elicited larger increases in class I HLA levels compared to PR8 infections. IAV genome segment 8 encodes the primary innate immune antagonist protein, non-structural protein 1 (NS1), which is highly variable between strains. The FM-MA NS1 protein lacks the carboxy-terminal 28 amino acids found in PR8 NS1 (Fig. 7A). To assess the role of NS1 in HLA upregulation, we infected A549 cells with FM-MA and PR8, as well as a panel of PR8 viruses with NS1 mutations that compromise its ability to suppress innate immune responses. These include point mutations in NS1 that disrupt its ability to suppress RIG-I activation (R38A, K41A or E96A, E97A) (40, 41) and a larger deletion that removes the effector domain and disordered carboxy-terminal tail (N80) (42). Consistent with known properties of NS1 in suppressing IFN production, all three NS1-mutant viruses caused HLA upregulation, and this upregulation was higher than the parental PR8 strain or the FM-MA strain (Fig. 7B, upper panel). Incubation of A549 cells with UV-inactivated culture supernatants from these infections revealed a key role for NS1 in limiting paracrine signalling. Conditioned media from FM-MA or PR8 infections caused moderate increases in HLA-A/B/C levels, whereas media from NS1 mutant virus infections elicited marked increases in HLA-A/B/C, and showed a trend towards increased HLA-B and HLA-E levels when measured independently (Fig. 7B, lower panel). Together, these findings clearly demonstrate that NS1 plays a lead role in suppressing the HLA presentation in IAV infected cells and bystander cells alike.

**Figure 7.**
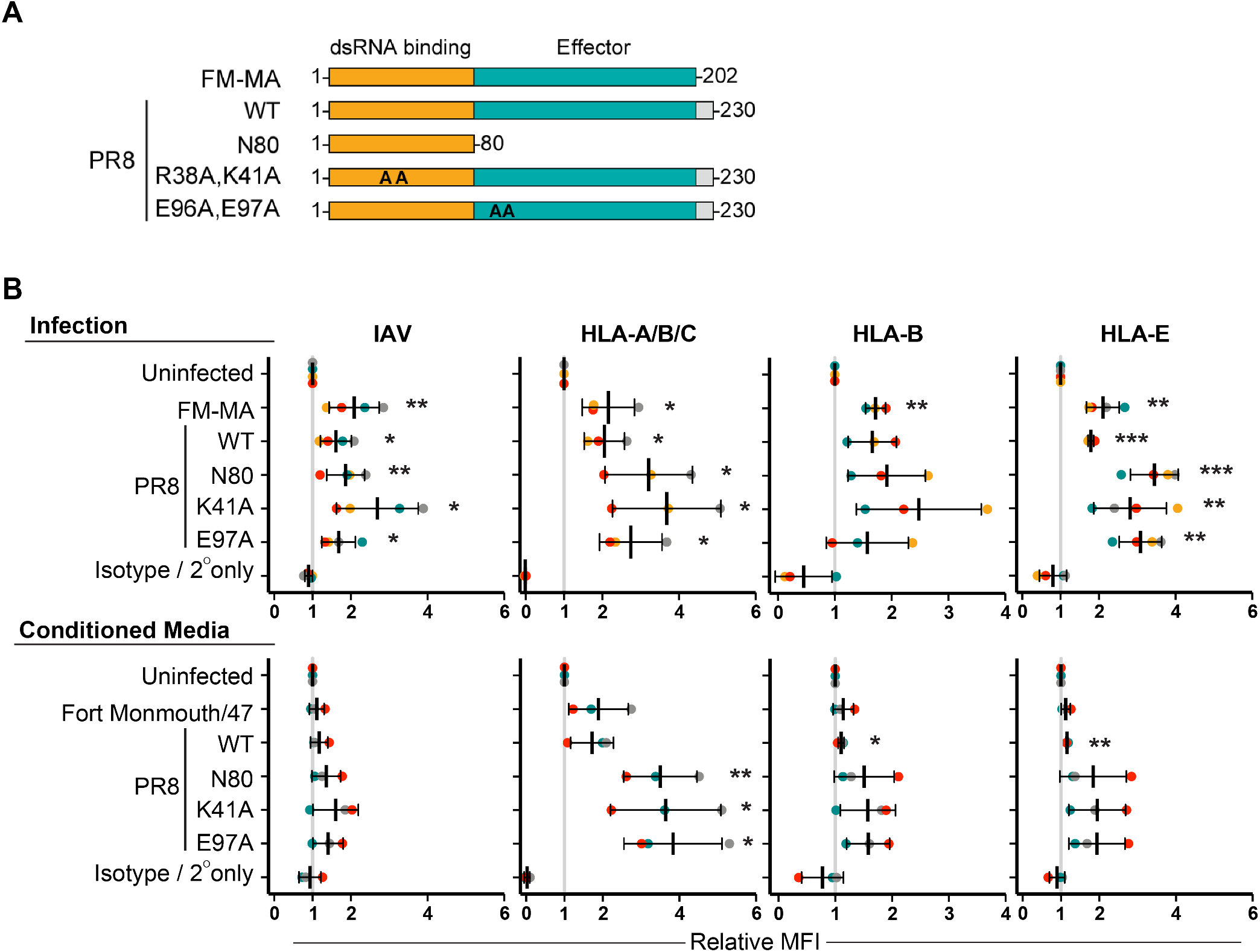
NS1 protein limits cell-intrinsic and paracrine upregulation of class I HLA proteins. (A) A diagram representing wild type and mutant NS1 proteins used in this study. A carboxy-terminal disordered ail region present in PR8 NS1 and absent in FM-MA NS1 is shown in grey. Positions of alanine substitutions n R38A,K41A and E96A,E97A mutant proteins are indicated as ‘AA’. Amino-terminal dsRNA binding domain is in orange; effector domain is in teal. (B) A549 cells were infected with the indicated viruses at an MOI=1 or mock-infected. At 17 hpi, cell supernatants were collected prior to cell fixation, and transferred to naïve A549 cells for an additional 17 h incubation prior to fixation. Donor and recipient cells were mmunostained with the indicated anti-HLA antibodies to determine cell surface levels of NK cell ligands; cells were subsequently permeabilized for immunostaining of intracellular IAV proteins and analyzed by flow cytometry. Top panels show data from donor infected or mock-infected cells. Bottom panels show data from cells exposed to conditioned media. Data is presented as MFI relative to uninfected cells or conditioned media treatment from uninfected cells. Each data point represents an independent experiment. Means and ±SD are shown. * p<0.05, ** p<0.01, *** p<0.001.

## DISCUSSION

NK cell receptors bind to ligands on the surface of infected cells and initiate antiviral immune responses by integrating activating and inhibitory signals. Here, we show that IAV infection of cultured epithelial cells alters expression of an array of ligands for activating and inhibitory NK cell receptors. With some exceptions, we observed a general trend towards increased expression of ligands for inhibitory receptors and downregulation of ligands for activating receptors, suggesting that the net effect of viral reprogramming of epithelial cells could be suppression of NK cell responses. Class I HLA proteins are recognized by KIR proteins on NK cells, and increased binding of KIR by HLA on IAV-infected cells *in vitro* has been shown to inhibit NK cell activity (28). However, the mechanisms that control HLA upregulation on IAV-infected cells are not fully understood. Here we report that class I HLA upregulation depends on post-entry steps in replication because UV-inactivated virus had no effect on HLA gene expression or accumulation of HLA proteins on the surface of A549 epithelial cells. We observed that defective viral RNAs produced during IAV replication were sufficient to induce expression and cell surface presentation of class I HLA on infected cells. Knowing that the RIG-I/MAVS signaling axis is the primary mechanism of detection of IAV replication intermediates in infected cells that drives antiviral responses, we tested HLA upregulation in MAVS deficient cells. We observed that genetic deletion of MAVS prevented class I HLA upregulation in response to IAV infection or ectopic expression of mvRNAs or DI RNAs, suggesting that aberrant viral RNAs generated during infection are bound by RIG-I and transduce signals that increase HLA gene expression.

Our work shows that IAV infection causes MAVS-dependent increases in expression of the antigen processing and presentation machinery including class I HLA-A, -B and -C and associated B2M, TAP1 and PSMB8 proteins, as well as the non-classical HLA-E protein. These comprise an antiviral gene expression program that responds to detection of defective viral RNAs by RIG-I. Cytotoxic T cells (CTL) and NK cells rely on HLA proteins for target cell recognition (43–45). Specifically, CTL activation and lysis of target cells requires binding to class I HLA proteins loaded with viral peptide antigens or HLA-E proteins loaded with noncanonical peptides from viruses and stress-related proteins (44, 46). By contrast, NK cell activation is inhibited by increased HLA protein levels on the surface of virus-infected cells by engaging inhibitory KIR proteins, as a main function of NK cells is to destroy host cells that have no surface expression of class I HLA proteins (28, 33, 34). Our observations are consistent with numerous reports of viruses that induce class I HLA expression or encode structurally similar immunoevasins that engage inhibitory receptors on NK cells and undermine their activity (29, 33, 34, 47–50). Thus, MAVS-dependent increases in cell surface class I HLA proteins have the potential to skew antiviral immune responses to thwart NK cells at the expense of potential CTL activation, which suggests that NK cells represent an existential threat for many viruses.

In the course of these studies we discovered that IFN can amplify responses to aberrant viral RNA products to increase HLA presentation. Specifically, we found that IAV infection of A549 cells stimulated production of IFN-β and IFN-λ1 in a MAVS-dependent manner, which dramatically increased at later times post-infection. Conditioned medium collected from these infected cells elicited modest, but significant, increases in HLA presentation on naïve epithelial cells that paled in comparison to the magnitude of increase on the donor infected cells. Conditioned medium collected from cells expressing IAV mvRNAs and DI RNAs similarly induced modest increases in surface class I HLA proteins when incubated with naïve A549 cells. We have not yet taken steps to fully characterize the composition of these culture supernatants, but the available evidence points to a role for type I IFNs, and, to a lesser extent, type III IFNs. However, it remains formally possible that additional factors secreted by infected cells could contribute to HLA gene expression.

HLA is upregulated in response to infection by a wide array of viruses, but underlying mechanisms differ. Hepatitis C virus (HCV) infection indirectly increases cell surface class I HLA levels by increasing expression of TAP1 and aiding transport of processed peptides to the endoplasmic reticulum where they can be loaded onto HLA and transported to the cell surface (34). Similarly, West Nile virus (WNV) infection increases TAP1 activity, resulting in increased transport of processed peptide antigens into the ER and higher levels of HLA:peptide complexes on the surface of infected cells (51). By contrast, Zika virus infection stimulates the RIG-I/MAVS/IRF3 pathway and downstream IFN-β expression, which increases HLA expression in infected cells (33). This mechanism is quite similar to the one we describe herein for IAV, except that in Zika virus infected cells RIG-I binds to the 5’-triphosphate end of the intact (+)-sense ssRNA virus genome (52) rather than defective RNA products of the IAV polymerase.

In this study we identified a viral protein, NS1, which normally prevents RIG-I-mediated detection of defective viral RNAs and downstream IFN signal transduction, that restrained class I HLA presentation. Indeed, NS1 not only suppressed HLA presentation on infected cells, but it also had a dramatic impact on HLA expression in bystander cells; treatment of naïve A549 cells with UV-inactivated culture supernatants collected from NS1 mutant virus infections elicited strong class I HLA upregulation compared to controls. However, NS1 has also been shown to increase transcription of the endoplasmic reticulum aminopeptidase 1 (*ERAP1*), which encodes a component of the antigen presentation machinery (53). Thus, the effect of NS1 on class I HLA-mediated antigen presentation is not limited to the effects mediated by IFN inhibition. More detailed studies of this hypervariable virulence factor will be required to fully understand the impact of NS1 on innate immune responses involving NK cells.

The existence of aberrant IAV RNA species has been well documented, but it has been less clear whether these products can benefit the virus. There is substantial evidence that defective RNA products of the viral polymerase limit productive viral replication by inducing IFN responses and promoting the generation of defective viral particles when incorporated into viral progeny. Our work demonstrates that aberrant IAV mvRNAs and DI RNAs stimulate class I HLA expression, which may aid viral evasion of NK cell-mediated immune responses.

## MATERIALS AND METHODS

### Cell lines

A549 cells and derivatives were cultured in complete Dulbecco’s modified Eagle’s medium (DMEM) supplemented with 10% fetal bovine serum (Thermo Fisher Scientific) at 37°C and 5% CO_2_. To generate A549-MAVS-KO cells, A549 cells were seeded at 1.65 × 10^5^ cells per well in a 12-well cluster dish to obtain a confluency of 80% the next day. One hour before transfection, medium was changed to F12 medium supplemented with 1% L-glutamine and 10% Fetalclone III serum (FCl-III) (Thermo Fisher Scientific). A total of 1.6 μg of Cas9 Nuclease Expression Plasmid (Dharmacon, #U-005200-120), 50 nM tracrRNA (Dharmacon) and 50 nM crRNA (crRNA non-targeting control 1 #U-007501-05 or crRNA human MAVS (Gene ID:57506) ex2, #GRANB-000259) were transfected with 40 μg/mL Dharmafect DUO transfection reagent (Dharmacon). 48 hours later, 2 μg/mL puromycin (Sigma Aldrich) was added to select for cells that have integrated the Cas9 expression plasmid. Cells were cultured for 7 days. Monoclonal populations were obtained by seeding cells at 40 cells/mL and isolation of clones using cloning rings. Gene editing was confirmed by Sanger sequencing at the Génome Québec Innovation Centre (McGill University, Montréal, QC). CRISP-ID web application tool (54) was used to locate the targeted region and monitor the insertions/deletions within the gene. crRNA non-targeting control sequence: GATACGTCGGTACCGGACCG. crRNA human MAVS (Gene ID: 57506) ex2 sequence: GGATTGGTGAGCGCATTAGA.

### Viruses and infections

Wild-type (WT) influenza A/Puerto Rico/8/1934 H1N1 (PR8) virus was generated using the 8-plasmid reverse genetic system (55) as previously described (40, 42). Viral stocks were produced in Vero cells and titers were determined by plaque assays in Vero cells. NS1 mutations were verified by Sanger sequencing of virus stocks. Mouse-adapted influenza A/Fort Monmouth/1/1947 (FM-MA) virus was a generous gift from Dr. Earl G. Brown (University of Ottawa). Viral stocks were produced in MDCK cells and infectious titers determined by plaque assays in MDCK cells. All plaque assays were performed using 1.2% Avicel overlays as described in Matrosovich *et al*. (56). Plaque assays and virus production in MDCK cells were performed in the presence of 1 μg/ml tosyl phenylalanyl chloromethyl ketone (TPCK)-treated trypsin (Sigma Aldrich), whereas similar procedures in Vero cells employed 2.5 μg/ml TPCK-treated trypsin. A549 cell monolayers were mock-infected or infected with the WT or mutant viruses at MOI=1 for 1 h at 37°C. Monolayers were washed with PBS and overlaid with fresh infection media (0.5% BSA in DMEM supplemented with 20 μM L-glutamine) and incubated at 37°C in 5% CO_2_ atmosphere.

### Plasmids, transfections and luciferase assays

Full-length, DI and mvRNA minireplicon plasmids were a generous gift from Dr. Aartjan te Velthuis (Cambridge University, Cambridge, UK). A549 cells were co-transfected using Lipofectamine 2000 (Thermo Fisher Scientific) with plasmids encoding the three polymerase subunits (PB1, PB2 and PA) and NP from A/Udorn/307/1972 (H3N2) IAV, a generous gift from Dr. Yoshihiro Kawaoka (University of Wisconsin-Madison), and luciferase reporter plasmids under the control of the interferon-stimulated response element (ISRE) promoter (firefly) and the CMV promoter (Renilla). At 24 h post-transfection, cells were washed with PBS and lysed in 1x Reporter Lysis Buffer (Promega). The dual luciferase assay was performed 24 h post-transfection using the Dual-Glo Luciferase Assay System (Promega).

### RNA purification, cDNA preparation and qPCR

RNA was extracted from cells and purified using the Quick-RNA miniprep kit (Zymo Research), following manufacturer’s protocol. In all cases, the RNA was treated with Turbo DNase (Life Technologies), then reverse transcribed using Verso cDNA synthesis kit (Thermo Fisher Scientific) according to the manufacturer’s protocol. qPCR was performed using iTaq Universal SYBR Green supermix (Bio-Rad) on a Bio-Rad CFX Connect instrument and analyzed using the Bio-Rad CFX Manager 3.1 program. Primers used are listed in Table 1.

**Table 1.**
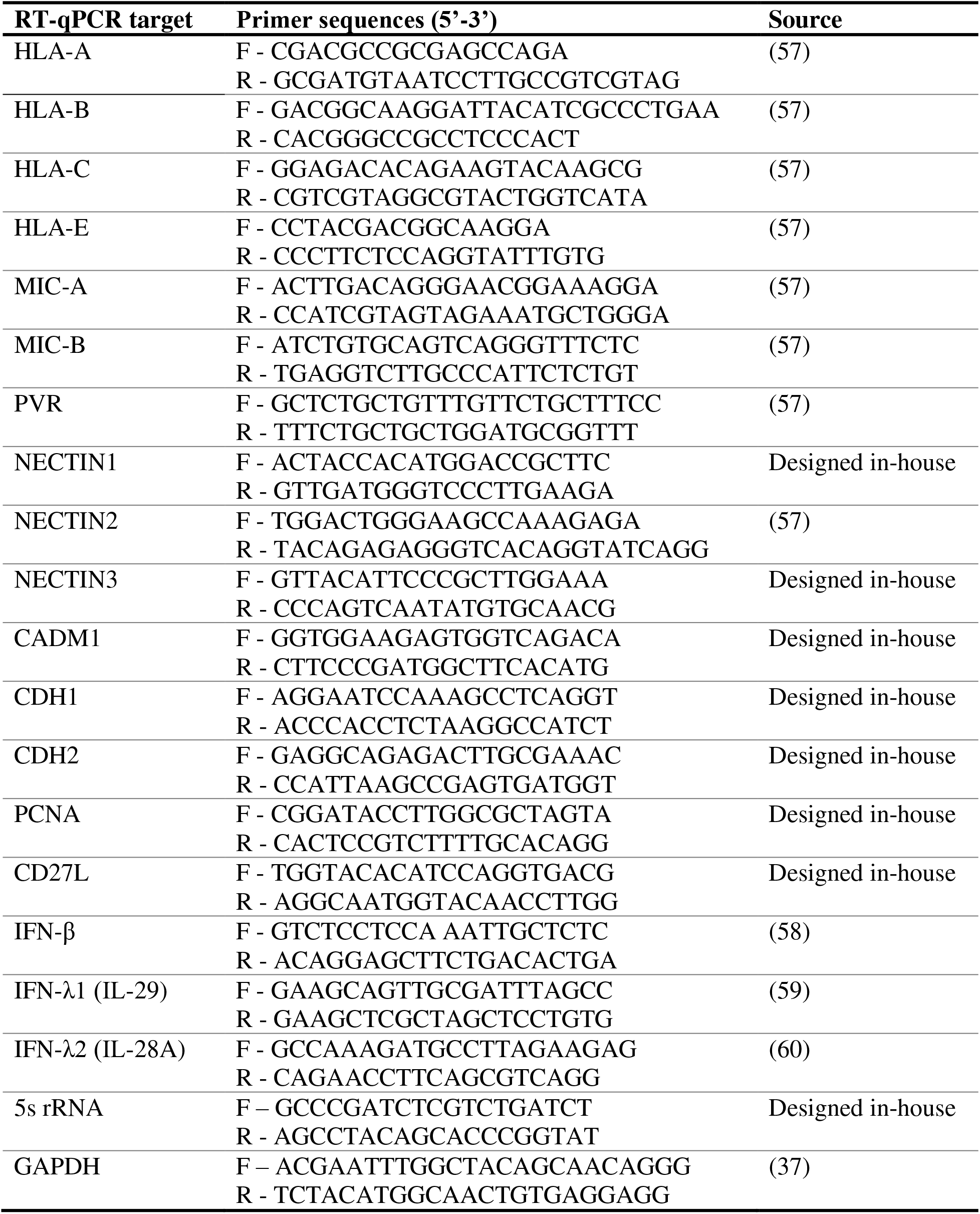
Primer sequences for RT-qPCR analysis

### Flow cytometry

IAV-infected, mock-infected, or transfected A549 cells or A549-MAVS-KO cells were resuspended using Versene solution (Thermo Fisher Scientific), washed with FACS buffer containing 0.5% BSA and 0.02% sodium azide in phosphate-buffered saline (PBS), and stained with fluorescently-conjugated antibodies against HLA-A/B/C (clone W6/32; BioLegend), HLA-B, HLA-E, CD155/PVR, CD113/NECTIN3, MICA/B antibodies in FACS buffer at 4℃ for 20 min. For intracellular detection of IAV proteins, cells were processed using fixation/permeabilization buffers (BioLegend) and stained with a goat polyclonal anti-IAV antibody (Abcam; ab20841). Transfected cells were fixed in 1% paraformaldehyde without permeabilization and intracellular staining. After a final wash in FACS buffer, cells were analyzed on a BD LSRFortessa FACS analyzer.

### Culture supernatant transfer experiments

Media from IAV-infected, mock-infected, or transfected A549 cells was collected and exposed to 1200 J/m^2^ UV light in a UV cross-linker to inactivate the virus. Naive A549 cells were treated with these culture supernatants for 17 h and cells were analyzed by flow cytometry as described above.

### Statistical analyses

Statistical significance for RT-qPCR and flow cytometry experiments were determined by two-way ANOVA with Sidak’s post-hoc test unless otherwise stated. For luciferase assays, statistical significance was determined by a one-way ANOVA test with Tukey post-hoc test and a cut-off *P* value of 0.05. * p<0.05, ** p<0.01, *** p<0.001, **** p<0.0001, and n.s., not significant.

## Acknowledgements

We thank members of the Makrigiannis and McCormick labs for critical reading of the manuscript. We thank Richard Webby (St. Jude Children’s Research Hospital), Aartjan te Velthuis (Cambridge University) and Earl G. Brown (University of Ottawa) for reagents. We thank Derek Rowter and Renee Raudonis in the Dalhousie University Flow Cytometry Core Facility for support. NZC was funded by a scholarship from Fonds de Recherche du Québec – Santé. This work was supported by Canadian Institutes for Health Research grants MOP-136817 (to C.M.), MOP-155906 (to A.P.M) and MOP-130527 (to NG) and by National Institutes of Health R01 AI137358 (to M.M.G.).

---

**Table S1.** List of publicly available gene expression data sets for bioinformatics analysis

| Accession # | IAV | Cell type* | Infection time (h) | MOI | Reference |
| --- | --- | --- | --- | --- | --- |
| GSE48466 | A/Ky/136/09 (Pdm) | NHBEC | 36 | 3 | (1) |
| GSE48466 | A/Ky/180/10 | NHBEC | 36 | 3 | (1) |
| GSE48466 | A/Brisbane/59/07 | NHBEC | 36 | 3 | (1) |
| GSE32138 | A/Udorn/307/72 | HAEC | 24 | 4 | (2) |
| GSE30723 | A/Puerto Rico/8/34 | ATII | 24 | 0.5 | (3) |
| GSE89008 | A/Vietnam/1203/04 | HTBE |  | 5 | (4) |
| GSE89008 | A/Wyoming/03/03 | HTBE | 18 | 5 | (4) |
| GSE89008 | A/California/04/09 | HTBE | 18 | 5 | (4) |
| GSE52930 | A/Puerto Rico/8/34(ΔNS1)<br>(vaccine backbone) | A549 | 12 | 3 | (5) |
| GSE48575 | A/Brisbane/59/07 | NHBEC | 24 | 0.9 | (6) |
| GSE48575 | A/Mexico/4108/2009 | NHBEC | 24 | 0.9 | (6) |
| GSE75699 | A/Puerto Rico/8/34 | NHBEC | 24 | 5 | (7) |
| GSE19392 | A/Puerto Rico/8/34 | NHBEC | 18 | 5 | (8) |
| GSE97949 | A/Anhui/1/2013 | A549 | 7 | 1 | (9) |
| GSE71766 | A/WS/33 | BEAS-2B | 12 | 2 | (10) |
| GSE36553 | A/Mexico/InDRE4487/2009 | A549 | 24 | 0.01 | (11) |
| GSE39200 | A/Texas/36/91 | HTBE | 9.5 | 2 | (12) |
| GSE31518 | A/Singapore/478/2009 | A549 | 10 | 4 | (13) |

\*NHBEC, normal human bronchial epithelial cells; HAEC, human airway epithelial cells; HTBE, human tracheobronchial epithelial cells; ATII, alveolar type II cells; A549, alveolar adenocarcinoma cell line; BEAS-2B, human bronchial epithelial cell line.

**Figure S1.**
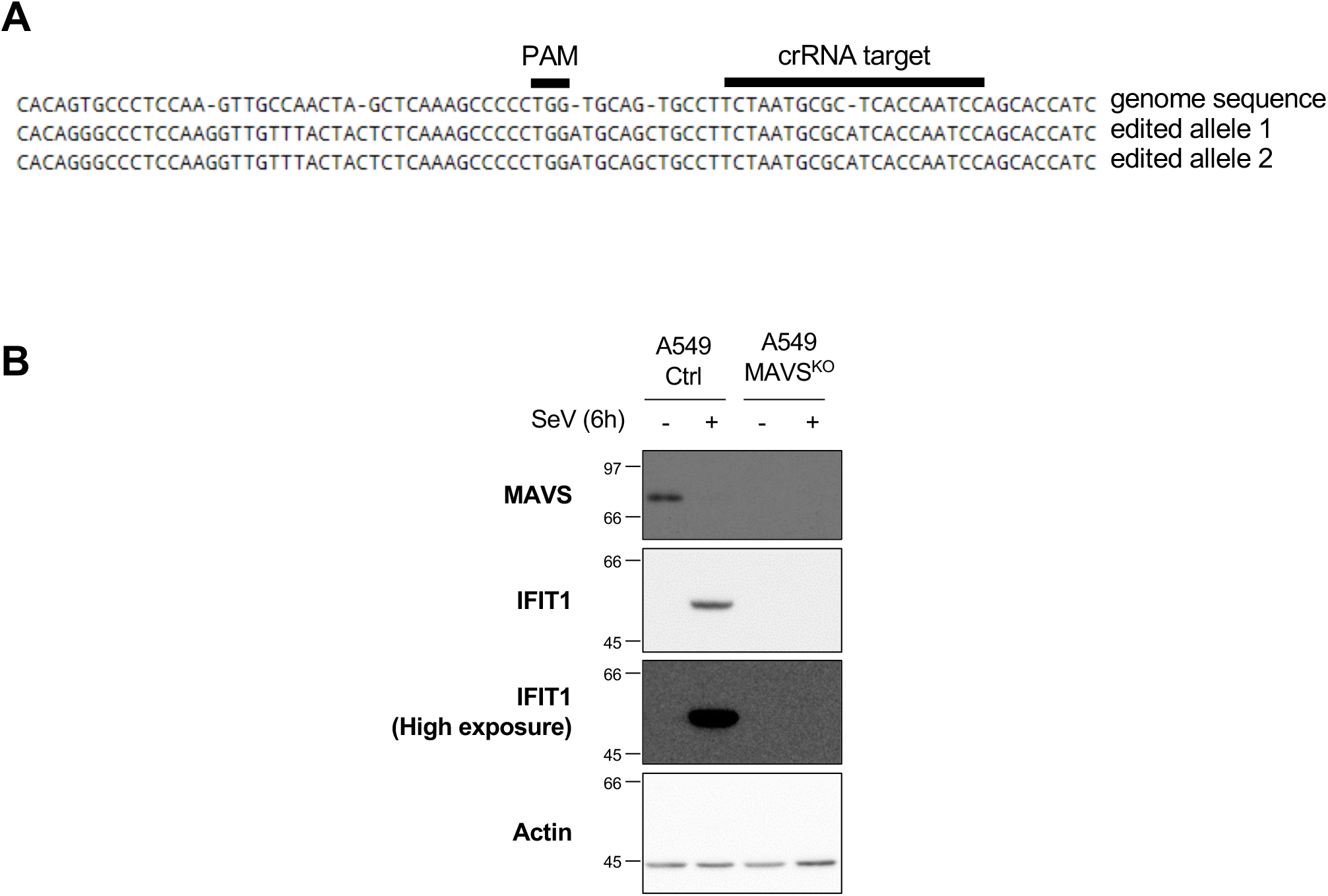
Validation of MAVS genome editing and impaired antiviral interferon responses. (A) A549 cells were subjected to CRISPR/Cas9 mediated cleavage using MAVS-specific CRSPR RNA (crRNA). Monoclonal cell populations were isolated, and disruption of the coding sequence was verified by Sanger sequencing: 2 insertions before and after the protospacer adjacent motif (PAM) and 1 insertion within the sequence targeted by the crRNA. CRISP ID web tool was used to analyze chromatograms (Dehairs, J. et al. CRISP-ID: decoding CRISPR mediated indels by Sanger sequencing Sci. Rep. 6, 28973; doi: 10.1038/srep28973 (2016). (B) A549 cells transfected with a non-targeting control crRNA (A549 ctrl) or a crRNA targeting MAVS genome (A549 MAVSKO) were infected or not with Sendai Virus (SeV) at 40HAU/106 cells for 6h. Whole Cell Extracts were resolved by standard SDS-PAGE and immunoblot. Proteins were detected using anti-actin, anti-IFIT1 and anti-MAVS. Left side of the gel: molecular weight (kDa) markers Note: at this time of SeV infection, MAVS is subjected to proteasome degradation.

